# A whole-genome scan for association with invasion success in the fruit fly *Drosophila suzukii* using contrasts of allele frequencies corrected for population structure

**DOI:** 10.1101/851303

**Authors:** Laure Olazcuaga, Anne Loiseau, Hugues Parrinello, Mathilde Paris, Antoine Fraimout, Christelle Guedot, Lauren M. Diepenbrock, Marc Kenis, Jinping Zhang, Xiao Chen, Nicolas Borowieck, Benoit Facon, Heidrun Vogt, Donald K. Price, Heiko Vogel, Benjamin Prud’homme, Arnaud Estoup, Mathieu Gautier

## Abstract

Evidence is accumulating that evolutionary changes are not only common during biological invasions but may also contribute directly to invasion success. The genomic basis of such changes is still largely unexplored. Yet, understanding the genomic response to invasion may help to predict the conditions under which invasiveness can be enhanced or suppressed. Here we characterized the genome response of the spotted wing drosophila *Drosophila suzukii* during the worldwide invasion of this pest insect species, by conducting a genome-wide association study to identify genes involved in adaptive processes during invasion. Genomic data from 22 population samples were analyzed to detect genetic variants associated with the status (invasive versus native) of the sampled populations based on a newly developed statistic, we called *C*_2_, that contrasts allele frequencies corrected for population structure. This new statistical framework has been implemented in an upgraded version of the program BayPass. We identified a relatively small set of single nucleotide polymorphisms (SNPs) that show a highly significant association with the invasive status of populations. In particular, two genes *RhoGEF64C* and *cpo*, the latter contributing to natural variation in several life-history traits (including diapause) in *Drosophila melanogaster*, contained SNPs significantly associated with the invasive status in the two separate main invasion routes of *D. suzukii*. Our methodological approaches can be applied to any other invasive species, and more generally to any evolutionary model for species characterized by non-equilibrium demographic conditions for which binary covariables of interest can be defined at the population level.

## Introduction

Managing and controlling introduced species require an understanding of the ecological and evolutionary processes that underlie invasions. Biological invasions are also of more general interest because they constitute natural experiments that allow investigation of evolutionary processes on contemporary timescales. Colonizers are known to experience differences in biotic interactions, climate, availability of resources, and disturbance regimes relative to populations in their native regions, often with opportunities for colonizers to evolve changes in resource allocation which favor their success (Balanya *et al.*, 2006; Dlugosch *et al.*, 2015; Lee and Gelembiuk, 2008). Adaptive evolutionary shifts in response to novel selection regimes may therefore be central to initial establishment and spread of invasive species after introduction (Colautti and Barrett, 2013; Colautti and Lau, 2015). In agreement with this adaptive evolutionary shift hypothesis, experimental evidence is accumulating that evolutionary changes are not only common during invasions but also may contribute directly to invasion success (Bock *et al.*, 2015; Colautti and Lau, 2015; Ellstrand and Schierenbeck, 2000; Facon *et al.*, 2011; Lee, 2002; Ochocki and Miller, 2017; Williams *et al.*, 2016). However, despite an increase in theoretical and empirical studies on the evolutionary biology of invasive species in the past decade, the genetic basis of evolutionary adaptations during invasions is still largely unexplored (Barrett, 2015; Reznick *et al.*, 2019; Welles and Dlugosch, 2018).

The spotted wing drosophila, *Drosophila suzukii*, represents an attractive biological model to study invasive processes. This pest species, native to South East Asia, initially invaded North America and Europe, simultaneously in 2008, and subsequently La Réunion Island (Indian Ocean) and South America, in 2013. Unlike most Drosophilids, this species lays eggs in unripe fruits by means of its sclerotized ovipositor. In agricultural areas, it causes dramatic losses in fruit production, with a yearly cost exceeding one billion euros worldwide (e.g., Asplen *et al.*, 2015; Cini *et al.*, 2012). The rapid spreading of *D. suzukii* in America and Europe suggests its remarkable ability to adapt or to acclimate to new environments and host plants. Using evolutionarily neutral molecular markers, Adrion *et al.* (2014) and Fraimout *et al.* (2017) finely deciphered the routes taken by *D. suzukii* in its invasion worldwide. Interestingly, both studies showed that North American (plus Brazil) and European (plus La Réunion Island) populations globally represent separate invasion routes, with different native source populations and multiple introduction events in both invaded regions (Fraimout *et al.*, 2017). These two major and separate invasion pathways provide the opportunity to evaluate replicate evolutionary trajectories. Finally, *D. suzukii* is a good model species for finely interpreting genomic signals of interest due to the availability of genome assemblies for this species (Chiu *et al.*, 2013; Ometto *et al.*, 2013; Paris *et al.*, 2019) along with the large amount of genomic and gene annotation resources available in its closely related model species *D. melanogaster* (Hoskins *et al.*, 2015).

In this context, advances in high-throughput sequencing technologies together with population genomics statistical methods offer novel opportunities to disentangle responses to selection from other forms of evolution. These advances are thus expected to provide insights into the genomic changes that might have contributed to the success in a new environment (reviewed in Bock *et al.*, 2015; Welles and Dlugosch, 2018). Hence, comparing the structuring of genetic diversity on a whole genome scale among invasive populations and their source populations might allow the characterization of the types of genetic variation involved in adaptation during invasion of new areas and their potential ecological functions. For example, Puzey and Vallejo-Marin (2014) used whole genome resequencing data to scan for shifts in site frequency spectra to detect positive selection in introduced populations of monkey-flower (*Mimulus guttatus*). Regions putatively under selection were associated with flowering time and abiotic and biotic stress tolerance and included regions associated with a chromosomal inversion polymorphism between the native and introduced range.

Identifying loci underlying invasion success can be considered in the context of whole-genome scan for association with population-specific covariate. These approaches, also known as Environmental Association Analysis (EAA), have received considerable attention in recent years (e.g., Coop *et al.*, 2010; de Villemereuil and Gaggiotti, 2015; Frichot *et al.*, 2013; Gautier, 2015). Most of the methodological developments have focused on properly accounting for the covariance structure among population allele frequencies that is due to the shared demographic history of the populations. This neutral covariance structure may indeed confound the relationship between the across population variation in allele frequencies and the covariates of interest (Coop *et al.*, 2010; Frichot *et al.*, 2013, 2015; Gautier, 2015). Yet, defining relevant environmental characteristics or traits as proxy for invasion success remains challenging and might even be viewed as the key aim. Therefore, we propose to simply summarize invasion success into a binary variable corresponding to the population’s historical status (i.e., invasive or native) based on previous studies. By extension, functional annotation of the associated variants identified may provide insights into candidate traits underlying invasion success (Estoup *et al.*, 2016; Li *et al.*, 2008; Wu *et al.*, 2019).

The Bayesian hierarchical model initially proposed by Coop *et al.* (2010), later extended in Gautier (2015) and implemented in the software BayPass, represents one of the most flexible and powerful frameworks to carry out EAA since it efficiently accounts for the correlation structure among allele frequencies in the sampled populations. Although association analyses may be carried out with categorical or binary covariables (see the example of *Littorina* population ecotypes in Gautier, 2015), the assumed linear relationship with allele frequencies is not entirely satisfactory and may even be problematic when dealing with small data sets or if one wishes to disregard some populations.

In the present study, we developed a non-parametric counterpart for the association model implemented in BayPass (Gautier, 2015). This new approach relies on a contrast statistic, we named *C*_2_, that compares the standardized population allele frequencies (i.e., the allele frequencies corrected for the population structure) between the two groups of populations specified by the binary covariable of interest. We evaluated the performance of this statistic on simulated data and used it to characterize the genome response of *D. suzukii* during its worldwide invasion. To that end, we generated Pool-Seq data (e.g., Gautier *et al.*, 2013; Schlotterer *et al.*, 2014) consisting of whole-genome sequences of pools of individual DNA (from n=50 to n=100 individuals per pool) representative of 22 worldwide populations sampled in both the invasive (n=16 populations) and native (n=6 populations) ranges of the species. We then estimated the *C*_2_ statistics associated with the invasive vs. native status of the populations on a worldwide scale or considering separately each of the two invasion routes (European and American) as characterized by Fraimout *et al.* (2017). Our aim was to identify genomic regions and genes involved in adaptive processes underlying the invasion success of *D. suzukii*.

### New Approaches

To identify single nucleotide polymorphisms (SNPs) associated with a population-specific binary trait, such as the invasive versus native status of *D. suzukii* populations, we developed a new statistic, we called *C*_2_. The *C*_2_ statistic was designed to contrast SNP allele frequencies between the two groups of populations specified by the binary trait while accounting for the possibly complex evolutionary history of the different populations. Indeed, the shared population history is a major (neutral) contributor to allele frequency differentiation across populations (e.g. Bonhomme *et al.*, 2010; Gunther and Coop, 2013) that may confound association signals (e.g. Coop *et al.*, 2010; Gautier, 2015).

We here relied on the multivariate normal approximation introduced by Coop *et al.* (2010) and further extended by Gautier (2015) to model population allele frequencies and to define the *C*_2_ contrast statistic. More precisely, consider a sample made of *J* populations (each with a label *j* = 1,…,*J*) that have been characterized for *I* bi-allelic SNPs (each with a label *i* = 1,…,*I*), with the reference allele arbitrarily defined (e.g., by randomly drawing the ancestral or the derived state). Let *α*_*ij*_ represent the (unobserved) allele frequency of the reference allele at SNP *i* in population *j*. As previously defined and discussed (Coop *et al.*, 2010; Gautier, 2015), we introduced an instrumental allele frequency 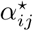 (for each SNP *i* and population *j*) taking values on the real line such that 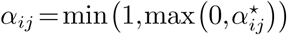).

Following Coop *et al.* (2010) and Gautier (2015), a multivariate Gaussian (prior) distribution of the vector 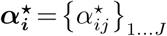 is then assumed for each SNP *i*:

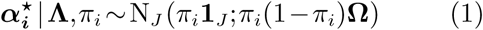

where **1**_*J*_ is the all-one vector of length *J*; **Ω** is the (scaled) covariance matrix of the population allele frequencies which captures information about their shared demographic history; and *π*_*i*_ is the weighted mean frequency of the SNP *i* reference allele. If **Ω** is used to build a tree or an admixture graph (Pickrell and Pritchard, 2012), *π*_*i*_ corresponds to the root allele frequency. We further define for each SNP *i* the vector 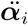 of standardized (instrumental) allele frequencies in the *J* populations as:

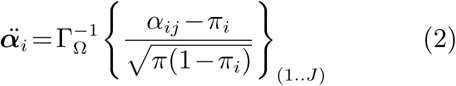

where Γ_Ω_ results from the Cholesky decomposition of Ω (i.e., Ω= Γ_Ω_^*t*^Γ_Ω_). The vector 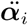 thus contains scaled allele frequencies that are corrected for both the population structure (summarized by **Ω**) and the across-population (e.g., ancestral) allele frequency (*π*_*i*_).

The *C*_2_ contrast statistic is then simply defined as the mean squared difference of the sum of standardized allele frequencies of the two groups of populations defined according to the binary trait modalities:

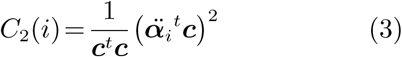

where 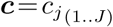 is a vector of the trait values observed for each population *j* such that *c*_*j*_ = 1 (respectively *c*_*j*_ = −1) if population *j* displays the first (respectively second) trait modality. One may also define *c*_*j*_ = 0 to exclude a given population *j* from the comparison.

According to our model, the *J* elements of 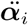 are independent and identically distributed as a standard Gaussian distribution under the null hypothesis of only neutral marker differentiation. The *C*_2_ statistic is thus expected to follow a *χ*^2^ distribution with one degree of freedom.

The estimation of the *C*_2_ statistic was performed here under the hierarchical Bayesian model implemented using a Markov-Chain Monte Carlo (MCMC) algorithm in the BayPass software (Gautier, 2015). However, such a multi-level modeling approach shrinks the estimated posterior means of the *C*_2_ toward their prior means, as already noticed in Gautier (2015) for the estimation of the SNP-specific *XtX* differentiation statistic defined as 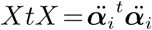 (Gunther and Coop, 2013). To ensure proper calibration of both the *C*_2_ and XtX estimates we thus relied on the scaled posterior means of the 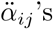, denoted 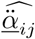 and computed as:

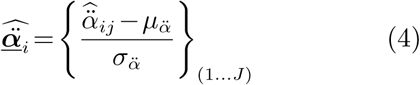

where 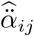 is the posterior means of 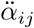 and 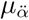 (respectively 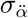) is the mean (respectively standard deviation) of the 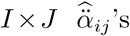 (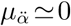 usually). The following estimators of XtX and *C*_2_, denoted for each SNP *i* as 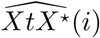 and 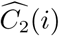 respectively, were then obtained as:

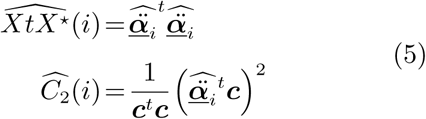

Under the null hypothesis, 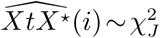 and 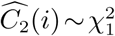 allowing one to rely on standard decision making procedures, e.g. based on p-values or more preferably on q-values to control for multiple-testing issues (Storey and Tibshirani, 2003).

## Results

### Simulation-based evaluation of the performance of our novel statistical framework

To evaluate the performances of the *C*_2_ contrast statistic for the identification of SNP associated with binary population-specific covariables, we simulated 100 data sets under the evolutionary scenario depicted in Figure 1A. Each simulated data set consisted of 5,000 SNPs genotyped for 320 individuals belonging to 16 differentiated populations subjected to two different contrasting environmental constraints, denoted *ec1* and *ec2* in Figure 1A. The *ec1* constraint was aimed at mimicking adaptation of eight pairs of geographically differentiated populations to two different ecotypes (e.g., host plant) replicated in different geographic areas. Conversely, the *ec2* might be viewed as replicated local adaptive constraints with a first type *a* specifying a large native area with several geographically differentiated populations (here six), and a second type *b* specifying invasive areas with differentiated populations originating from various regions of the native area (i.e., not related to the same extent to their contemporary native populations). It should be noted that the two *ec1* types were evenly distributed in the population tree while for *ec2*, the type *b* was over-represented in 10 populations (Figure 1A). During the adaptive phase, the fitness of individuals in the environment of their population of origin was determined by their genotypes at 25 SNPs for *ec1* and 25 SNPs for *ec2* constraints (hereafter referred to as *ec1* and *ec2* selected SNPs, respectively). Overall, the realized *F*_*ST*_ (Weir and Cockerham, 1984) ranged from 0.110 to 0.122 (0.116 on average) across the data sets, a level of differentiation similar to that observed in our worldwide *D. suzukii* sample (see below).

**FIG. 1.**
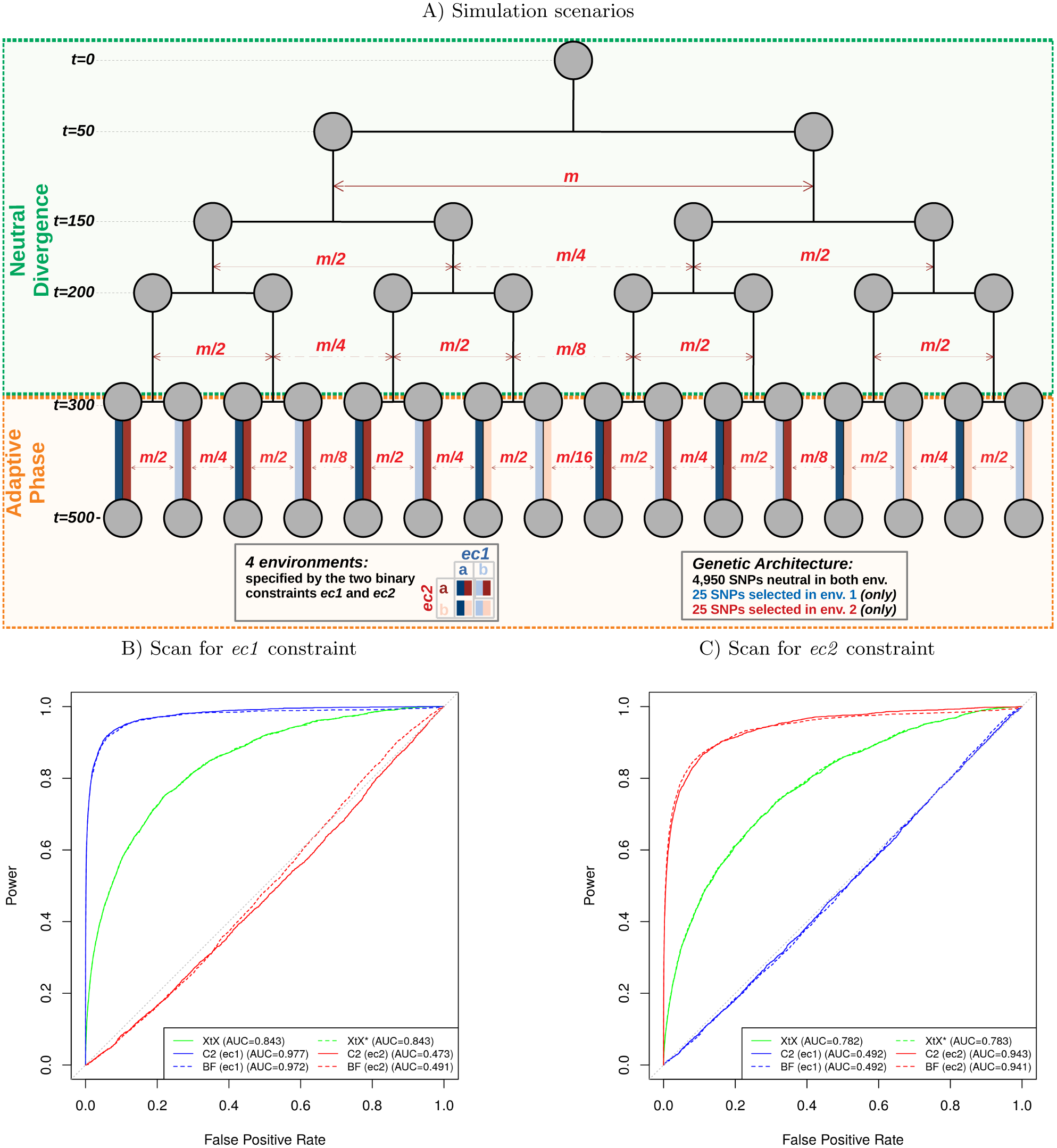
Evaluation of the performance of the *C*_2_ contrast statistic on simulated data and comparison with the BF for association and two XtX SNP-specific differentiation estimators. A) Schematic representation of the demographic scenario used for the simulation. It consists of two successive phases: (i) a neutral divergence phase with migration (only some illustrative migration combinations being represented) leading to the differentiation of an ancestral population into 16 populations after four successive fission events (at generations *t* = 50, *t* = 150, *t* = 200 and *t* = 300); and (ii) an adaptive phase (lasting 200 generations) during which individuals were subjected to selective pressures exerted by two environmental constraints (*ec1* and *ec2*) each having two possible modalities (*a* or *b*) according to their population of origin (i.e., eight possible environments in total). Out of the 5,000 simulated SNPs, the fitness of individuals in the environment of their population of origin was determined by their genotypes at 25 SNPs for *ec1* and 25 SNPs for *ec2* constraints. In total 100 data sets were simulated. B) and C) The ROC curves associated to the *ec1* and *ec2 C*_2_ contrasts and the two corresponding BF for association are plotted together with those associated with the two XtX estimators (i.e., posterior mean estimator XtX, and the new calibrated estimator *XtX*^*\**^). The FPR’s associated to each statistic were obtained from the corresponding neutral SNP estimates combined over the 100 simulated data sets (*n* = 4,950 × 100 = 495,000 values in total). Similarly, the TPR’s were estimated from either the *n* = 2,500 combined *ec1* (B) or *ec2* (C) selected SNPs. ROC AUC values are given between parentheses.

We further estimated with BayPass (Gautier, 2015) the *C*_2_ statistics for each *ec1* or *ec2* contrasting environmental constraints together with the corresponding Bayes Factors (BF) as an alternative measure of the support for association. For comparison purposes, we also estimated the SNP XtX differentiation statistic, using both the posterior mean estimator (Gautier, 2015) and the 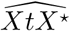 estimator described above. Note however that, as an overall (covariate-free) differentiation statistic, the XtX does not distinguish outlier SNPs responding to the *ec1* constraint from those responding to the *ec2* constraint.

Based on the status of each simulated SNPs (i.e., neutral, and *ec1* or *ec2* selected) and combining results in the 100 simulated data sets, standard receiver operating curves (ROCs) were computed (Grau *et al.*, 2015) and plotted in Figure 1B (respectively 1C) for the six statistics. This allowed comparing for various thresholds covering their range of variation of the different statistics, the power to detect *ec1* (respectively *ec2*) selected SNPs (i.e., the proportion of true positives among the corresponding selected SNPs) as a function of the false positive rates (FPR, i.e., the proportion of positives among neutral SNPs). The *C*_2_ statistic was found efficient to detect SNPs affected by *ec1* and *ec2* environmental constraints, the area under the ROC curve (AUC) being equal to 0.977 (Figure 1B) and 0.943 (Figure 1C), respectively. The unbalanced population representation of the two *ec2* types had a limited impact on the performance of the *C*_2_ statistic to identify the underlying selected SNPs. In addition, the *C*_2_ statistics clearly discriminated the selected SNPs according to their underlying environmental constraint. In other words, no selection signal was identified by the *C*_2_ statistic computed for the *ec2* (respectively *ec1*) contrast on *ec1* (respectively *ec2*) selected SNPs, resulting in ROC AUC close to the value of 0.5 obtained with a random classifier.

The ROC curves displayed in Figures 1B and 1C also revealed nearly identical performance of the *C*_2_ statistic and the BF. Accordingly, the correlation between both statistics were fairly high (Pearson’s *r* equal to 0.983 and 0.923 for *ec1* and *ec2*, respectively). Yet, one practical advantage of the *C*_2_ statistic was its good calibration with respect to the null hypothesis of no association, the corresponding p-values (assuming a *χ*^2^ distribution with 1 degree of freedom) being close to uniform (Figure S1).

Similarly, the two XtX estimators were found highly correlated (Pearson’s *r* = 0.998) with almost confounded ROC curves, but only the 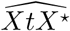 was properly calibrated (Figure S2). Their performances were however clearly worse than those obtained with the *C*_2_ (and BF) statistics. This was in part explained by their inability to discriminate between the two types of selected SNPs, selected SNPs overly differentiated in *ec2* generating false positives in the identification of *ec1* SNPs (Figure 1B) and vice versa. Accordingly, ROC AUC in Figure 1B for the XtX were also smaller than in Figure 1C, *ec1* selected SNPs being more differentiated than those in *ec2* due to the simulated design. Yet, the power of the XtX statistic to detect *ec1* or *ec2* selected SNPs remained substantially smaller than that of the corresponding *C*_2_ contrast statistics. For instance, at the 1% p-value significance threshold, the power to detect ec1 (respectively ec2) selected SNPs was equal to 72.6% (respectively 59.1%) with the *C*_2_ statistic and only 17.1% (respectively 10.4%) with the 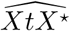 estimator, even when considering for the latter, a unilateral test to only target overly differentiated SNPs. Note that, as expected from the good calibration of the 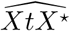 statistic, similar results were obtained when considering empirical p–value thresholds computed from the distribution of the XtX statistics estimated from neutral SNPs.

### Genome-wide scan for association with invasion success in *D. suzukii*

To identify genomic regions associated with the invasion success of *D. suzukii*, we carried out a genome scan, based on the *C*_2_ statistic, to contrast the patterns of genetic diversity among 22 populations originating from either the native (n=6 populations) or invaded areas (n=16 populations) (Figure 2A). To that end we sequenced pools of 50 to 100 individuals representative of each population (Table S1) and mapped the resulting sequencing reads onto the newly released WT3-2.0 *D. suzukii* genome assembly (Paris *et al.*, 2019). These Pool-Seq data allowed the characterization of 11,564,472 autosomal and 1,966,184 X–linked SNPs segregating in the 22 populations that were sub-sampled into 154 autosomal and 26 X–linked data sets (of ca. 75,000 SNPs each) for further analyses.

**FIG. 2.**
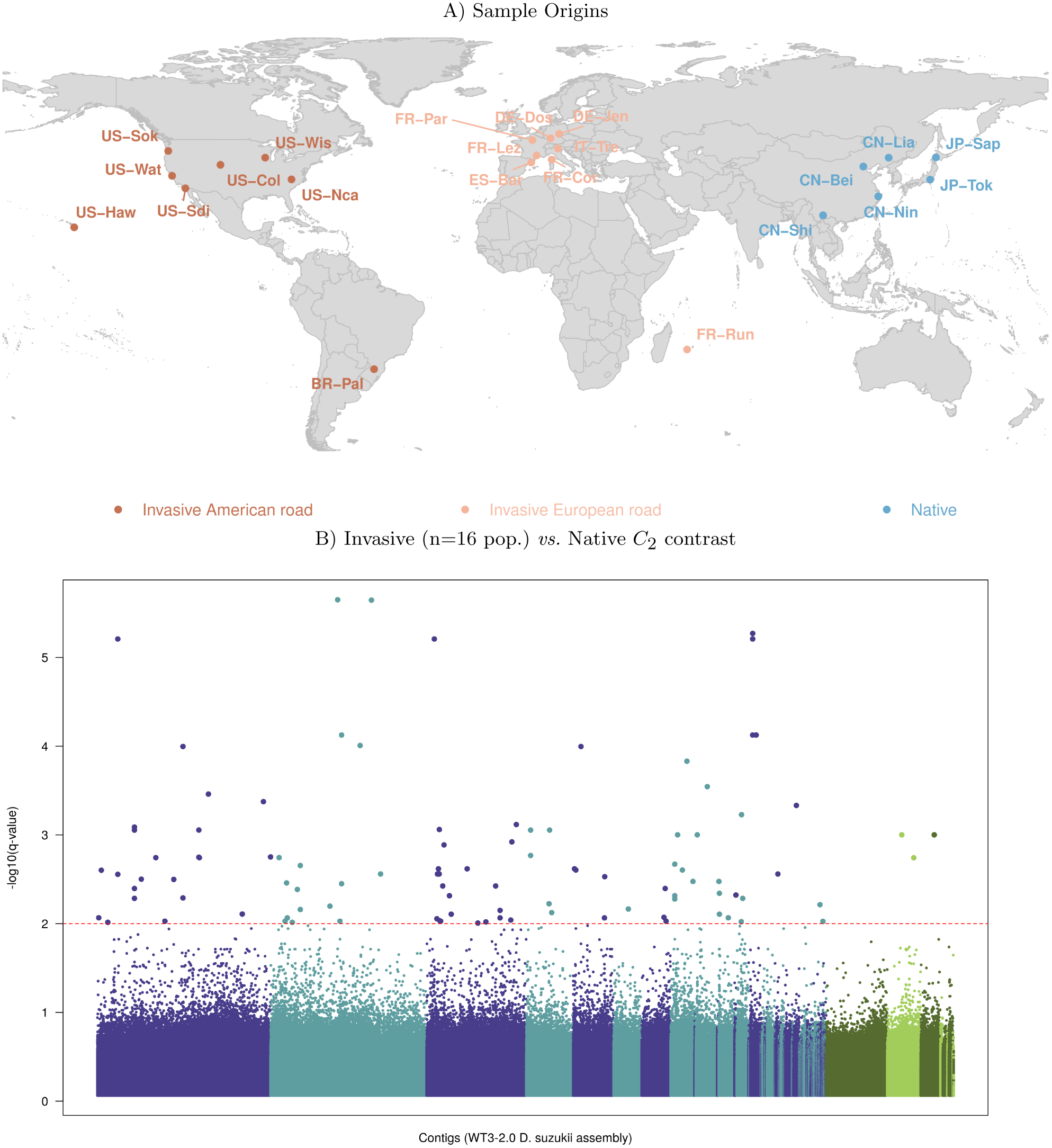
Whole-genome scan for association with invasion success in *D. suzukii*. A) Geographic location of the 22 *D. suzukii* population samples genotyped using a pool-sequencing methodology. Population samples from the native range are in blue and those from the invasive range are in red (American invasion route) or light red (European invasion route) (Fraimout *et al.*, 2017). See Table S1 for details on each population sample. B) Manhattan plot of the SNP q-values on a −log_10_ scale derived from the estimated *C*_2_ statistics for the native *vs.* invasive status contrast of the 22 worldwide *D. suzukii* populations. SNPs are ordered by their position on their contig of origin displayed with alternating dark blue and light blue color when autosomal and dark green and light green when X–linked. The horizontal dashed line indicates the 1% q-value threshold (here corresponding to a p-value threshold of 8.49 × 10^−8^) which gives the expected FDR (False Discovery Rate), i.e., the expected proportion of false positives among the 110 SNPs (highlighted in the plot) above this threshold.

The overall differentiation was estimated using the recently developed *F*_*ST*_ estimator for Pool-Seq data (Hivert *et al.*, 2018). It ranged from 8.86% to 9.02% (8.95% on average) for the autosomal data sets and from 17.6% to 17.8% (17.8% on average) for the X–chromosome data sets. Although a higher genetic differentiation is expected for the X-chromosome even under equal contribution of males and females to demography, the almost twice higher overall differentiation observed for the X chromosome compared to autosomes might have been accentuated by unbalanced sex-ratio (e.g., polyandry), male–biased dispersal or a higher impact of selection on the X–chromosome (Clemente *et al.*, 2018). Inferring sex-specific demography was beyond the scope of the present study, but for our purposes, this finding justified to perform separate genome scans on autosomal and X–linked SNPs.

We ran BayPass on the different data sets to estimate, for every SNPs, the *C*_2_ statistic that contrasts the allele frequencies of native and invasive populations, while accounting for their shared population history as summarized in the scaled covariance matrix **Ω**. Interestingly, the estimated **Ω** matrices for autosomal and X– linked SNPs resulted in a similar structuring of the genetic diversity across the 22 populations (Figure S3), which may rule out selective forces as the main driver of the differences of global differentiation levels observed between the two chromosome types. As expected from the simulation results, the distribution of the p-values derived from the *C*_2_ statistics was well-behaved, being close to uniform for higher p–values (Figure S4A). To account for multiple testing issues, we used the *qvalue* R package (Storey and Tibshirani, 2003) to compute the individual SNP q–values plotted in Figure 2B.

A striking feature of the resulting Manhattan plot was the lack of clustering of SNPs with high q–values which might be related to a small extent of linkage disequilibrium (LD) across the *D. suzukii* populations, as expected from their large effective populations sizes (Fraimout *et al.*, 2017). We identified 101 SNPs (including three X–linked) that were significant at the 1% q– value threshold (i.e., 1% of these 101 SNPs are expected to be false positives). As a matter of comparison, we also estimated the BF for association of the (standardized) population allele frequencies with the native or invasive status of the population, i.e., under a parametric regression model (Gautier, 2015) (Figure S5A). Out of the 101 significant SNPs previously identified, 80 displayed a BF> 20 db, the threshold for decisive evidence according to the Jeffreys’ rule (Jeffreys, 1961). However, in total, 6,406 SNPs displayed a BF> 20 db probably as a consequence of these BF’s not accounting for multiple testing issue. We also compared the *C*_2_ statistic to the *XtX* measure of overall differentiation. The (two-sided) p–values derived from the latter were also well behaved (Figure S4B) and allowed the computation of q–values to control for multiple testing. As shown in Figure S5B, at the same 1% q–value threshold for *XtX*, 71 out of the 101 *C*_2_ significant SNPs were significantly differentiated but they represented only a small proportion of the 35,546 significantly differentiated SNPs. This is not surprising since invasion success is obviously not the only selective constraint exerted on the 22 worldwide populations considered here.

The North-American (plus Brazil) and European (plus La Réunion Island) populations globally represent separate invasion routes that can be considered as two independent invasive replicates (Figure 2A). Interestingly enough, this feature of historical invasion fits well with the overall pattern of structuring of genetic diversity inferred from the **Ω** matrix estimated with our Pool-Seq data (see above and Figure S3). To identify signals common or specific to each invasion routes, we estimated the *C*_2_ statistic associated with the invasive vs. native status focusing either on the native and invasive populations of the European invasion route 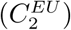, or native and invasive populations of the American invasion route 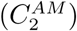. Note that the two invasion routes were both represented by eight invasive populations, suggesting similar power for the two 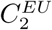 and 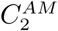 statistics. As observed above, the distribution of p–values derived from 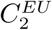 and 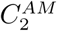 were found well behaved (Figures S4C and S4D, respectively) and hence q–values to control for multiple testing could be confidently computed. The cross-comparisons of the *C*_2_ statistics considering the 22 worldwide populations (hereafter denoted 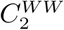), the 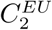 and the 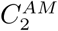 are plotted in Figures 3A (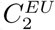 versus 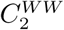), 3B (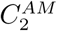 versus 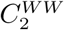) and 3C (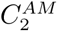 versus 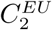).

**FIG. 3.**
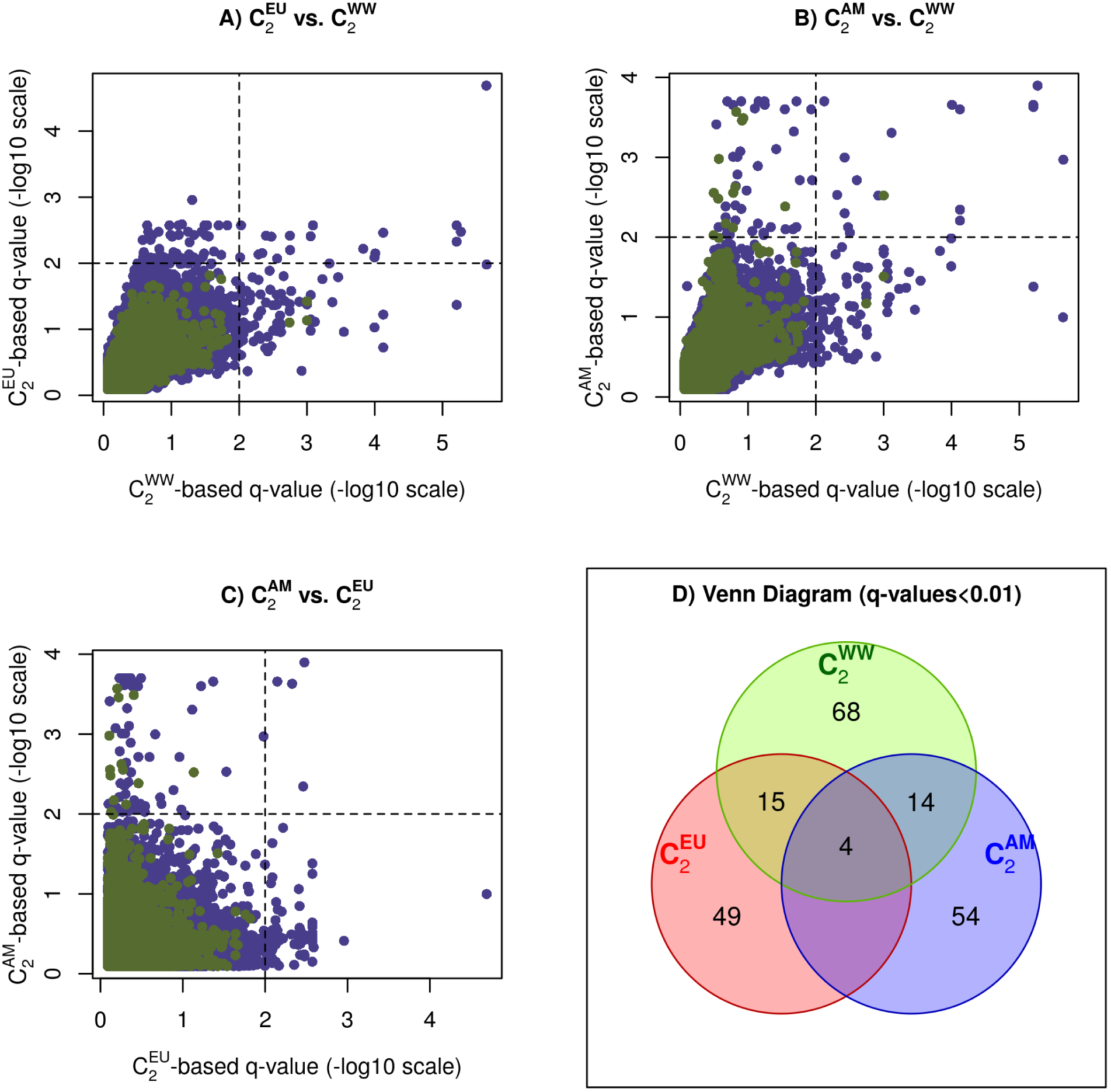
Pairwise comparison of the q–values derived from the 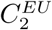 (native vs. invasive *D. suzuki* populations of the European invasion route) versus the 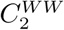 (native vs. worldwide invasive populations) statistics A), the 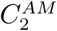 (native vs. invasive populations of the American invasion route) versus the 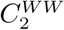 statistic B), and the 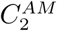 versus the 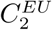 statistics C). In A), B) and C) the dashed vertical and horizontal lines indicate the 1% q–value threshold for the *C*_2_ derived q–values. D) Venn diagram of the number of SNPs significant at the 1% q–values among the three contrast analyses (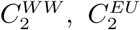 and 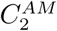). Values for the autosomal (X–linked) SNPs are plotted in purple (green).

In total, 204 SNPs (detailed in Table S2) were significant in at least one of the three contrasts at the 1% q–value threshold. The overlap among the three different sets of significant SNPs was summarized in the Venn diagram displayed in Figure 3D. Among the 68 SNPs significant for the 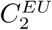, 15 were also significant for 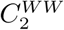 and 49 were not significant in the other tests. Likewise, among the 72 SNPs found significant for the 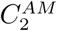, 14 were also significant for 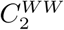 and 54 were not significant in the other tests. Hence, the majority of the significant SNPs identified with either the 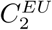 or the 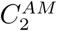 contrasts might be viewed as specific to one of the two invasion routes, the signal being lost in the global worldwide comparison for a substantial proportion of them. This is presumably due to a reduced power resulting from the addition of non-informative populations when computing the 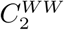 statistic. Conversely, 68 SNPs found significant with 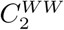 were neither significant with 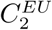 nor 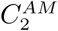 contrasts. These SNPs might correspond to partially convergent signals among the two invasion routes (i.e., the informative populations are distributed among the two routes). Most interestingly, four SNPs were found significant at the 1% q–value threshold in the three contrast analyses (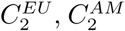 and 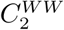) and might thus be viewed as strong candidates for association with the global worldwide invasion success of *D. suzukii*.

### Annotation of candidate SNPs

For annotation purposes, we relied on genomic resources available in *D. melanogaster*, a model species closely related to *D. suzukii*. More specifically we extracted from the WT3-2.0 *D. suzukii* genome assembly 5 kb long genomic sequences surrounding each of the 204 SNPs identified above and aligned them onto the *dmel6* reference genome (Hoskins *et al.*, 2015) using the BLAT algorithm implemented in the program *pblat* (Wang and Kong, 2019). The gene annotation available from the UCSC genome browser allowed us to map 169 SNPs out of the 204 SNPs onto 130 different *D. melanogaster* genes, 145 SNPs lying within the gene sequences and 24 less than 2.5 kb apart (our predefined threshold; Table S2). Only one of the four SNPs significant for the three contrasts (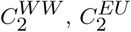 and 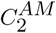) could not be assigned to a *D. melanogaster* gene, because its derived 5 kb long sequences aligned onto a *D. melanogaster* sequence located 10 kb away from the closest annotated gene.

Most of the 130 identified genes (80%) were represented by a single SNP, a feature in agreement with the visual lack of clustering of SNPs with strong signal already observed in the Manhattan plot (Figure 2B). It should be noticed that 14 of the 130 genes (ca. 11%) were long non-coding RNA. We however decided to focus on the 26 genes that were represented by at least two SNPs significant in one of the three contrast analyses; see Table 1 for details. The significant SNPs underlying the different genes tended to be very close, spanning a few bp (span > 1kb for only five genes). In particular, we observed doublet variants (i.e., adjacent SNPs in complete LD) within three genes (*cpo, ome* and *lnc:CR45759*).

**Table 1.** Description of the 26 orthologous *D. melanogaster* genes represented by at least two of the 204 SNPs found significant for one of the three contrast analyses, 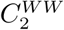 (6 native *vs.* 16 invasive populations), 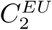 (6 native *vs.* 8 invasive populations of the European invasion route) and 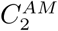 (6 native vs. 8 invasive populations of the American invasion route). The third column gives the overall number of significant SNPs (at the 1% q–value threshold) and their maximal spacing in bp (on the *D. suzukii* assembly). Columns 4 to 6 gives the number of significant SNPs for each of the three contrast analyses.

| *D. melanogaster* Gene<br>(Full Name) | Position on *dmel6*<br>(in kb) | Number of significant SNPs |  |  |  |
| --- | --- | --- | --- | --- | --- |
| | | All $C_2$ (dist. in bp) | $C_2^{WW}$ | $C_2^{EU}$ | $C_2^{AM}$ |
| Der-1 (Derlin-1) | chr2L:1,974-1,975 | 2 (236) | 1 | - | 1 |
| Gdi (GDP dissociation inhibitor) | chr2L:9,492-9,495 | 4 (342) | 4 | 4 | - |
| lncRNA:CR45693 (long non-coding RNA) | chr2L:14,51-14,512 | 2 (14) | 2 | 1 | - |
| Tpr2 (tetratricopeptide repeat protein 2) | chr2L:16,492-16,507 | 2 (8) | - | 2 | - |
| Ret (Ret oncogene) | chr2L:21,182-21,199 | 2 (70) | 2 | - | - |
| tou (toutatis) | chr2R:11,579-11,616 | 2 (18) | 1 | - | 2 |
| jeb (jelly belly) | chr2R:12,091-12,119 | 2 (14) | 2 | - | - |
| CG5065 | chr2R:16,608-16,625 | 2 (13) | - | 2 | - |
| bab2 (bric a brac 2) | chr3L:1,140-1,177 | 2 (11189) | 1 | - | 1 |
| axo (axotactin) | chr3L:4,630-4,687 | 2 (25886) | - | 1 | 1 |
| RhoGEF64C ($\rho$ guanine nucl. exch. fact. at 64C) | chr3L:4,693-4,796 | 2 (8) | 2 | 1 | 1 |
| CG7509 | chr3L:4,803-4,805 | 2 (5) | - | 2 | - |
| Con (connectin) | chr3L:4,938-4,976 | 2 (616) | 1 | 1 | - |
| Ets65A (Ets at 65A) | chr3L:6,098-6,124 | 2 (27998) | 1 | 1 | - |
| lncRNA:CR45759 (long non-coding RNA) | chr3L:6,787-6,787 | 4 (106) | - | - | 4 |
| ome (omega) | chr3L:14,673-14,748 | 2 (1) | 2 | - | - |
| sa (spermatocyte arrest) | chr3L:21,405-21,407 | 2 (61) | 1 | 1 | - |
| yellow-e (yellow-e) | chr3R:13,410-13,415 | 3 (33) | 3 | - | 1 |
| cv-c (crossveinless c) | chr3R:14,392-14,482 | 4 (2737) | 1 | - | 3 |
| osa (osa) | chr3R:17,688-17,718 | 2 (29) | - | - | 2 |
| cpo (couch potato) | chr3R:17,944-18,016 | 3 (193) | 3 | 2 | 3 |
| Rh3 (rhodopsin 3) | chr3R:20,081-20,082 | 2 (5709) | 2 | 1 | - |
| Ctl2 (choline transporter-like 2) | chr3R:29,123-29,128 | 2 (3) | - | - | 2 |
| Syt12 (synaptotagmin 12) | chrX:13,359-13,368 | 3 (65) | 1 | - | 2 |
| Ac13E (adenylyl cyclase 13E) | chrX:15,511-15,554 | 4 (19) | - | - | 4 |
| Axs (abnormal X segregation) | chrX:16,680-16,684 | 2 (11) | - | - | 2 |

Among these 26 candidate genes, 10 and 12 might be considered as specific to the European and American invasion routes, respectively, since they did not contain any SNP significant for the alternative contrasts. Only two genes contained SNPs significant in all three contrast analyses: *RhoGEF64C* with one SNP and *cpo* with two SNPs. Such convergent signals of association with invasive status in the two independent invasion routes were particularly convincing. The median allele frequencies (computed from raw read counts) for the reference allele underlying the corresponding *RhoGEF64C* significant SNP was 0.09 (from 0.00 to 0.44) in the native populations compared to 0.93 (from 0.90 to 0.98) and 0.87 (from 0.59 to 1.00) in the invasive populations of the European and American invasion routes, respectively (Table S2). Similarly, the two SNPs significant for the three contrast analyses in the *cpo* gene actually formed a doublet with a median reference allele frequency of 0.20 (from 0.02 to 0.33) in the native populations compared to 0.99 (from 0.91 to 1.00, excluding the outlying Hawaiian population) in the invasive populations of the European and American invasion routes, respectively (Table S2). Finally, for both the genes *RhoGEF64C* and *cpo*, all *D. suzukii* extended sequences underlying the corresponding SNPs aligned within potentially rapidly evolving intronic sequences. These sequences nevertheless displayed substantial similarities with other related drosophila species, as shown in Figure S6 for the gene *cpo*.

## Discussion

We characterized the genome response of *D. suzukii* during its worldwide invasion by conducting a genome-wide scan for association with the invasive or native status of the sampled populations. To that end, we relied on the newly developed *C*_2_ statistic that was aimed at identifying significant allele frequencies differences between two contrasting groups of populations while accounting for their overall correlation structure due to the shared population history. Our approach identified genomic regions and candidate genes most likely involved in adaptive processes underlying the invasion success of *D. suzukii*.

Overall, we found that a relatively small number of SNPs were significantly associated with the invasive status of *D. suzukii* populations. This may seem surprising since the binary trait under study (invasive versus native) is complex in the sense that numerous biological differences may characterize invasive and native populations. The invasion process itself, including the associated selective pressures and the genetic composition of the source populations, may actually differ depending on the considered invaded areas. Hence the small number of SNPs showing strong signals of association with the invasive status may stem from the integrative nature of our analysis over a large number of invasive populations from different invasion routes. The genomic features that may be identified under this evolutionary configuration are expected to correspond to major genetic changes instrumental to invasions shared by a majority of populations. Accordingly, it is worth noting that the independent contrast analyses of the two main invasion routes (i.e. the American and the European routes) point to substantially different subsets of SNPs significantly associated with the invasive status of the populations. This suggests that the source populations and some aspects of the invasion process differ in the two invaded areas. This could however also reflect the presumably polygenic nature of the traits underlying invasion success since the evolutionary trajectories of complex traits may rely on different combination of favorable genetic variants.

The availability of a high quality genome assembly of *D. suzukii* (Paris *et al.*, 2019) and a large amount of genomic resources for its sister model species *D. melanogaster* allowed identifying a set of genes associated with the invasive status of populations. A subset of those genes was associated with physiological functions and traits previously documented in *D. melanogaster*, but for most of them, functional and phenotypic studies turned out to be limited. Their putative role in explaining the invasion success thus remained largely elusive. To avoid too speculative interpretations (Pavlidis *et al.*, 2012), we will not elaborate further on the candidate genes. Yet, we did notice that long non–coding RNAs represent more than 10% (14 out of 130) of our candidate genes, a feature which may underline a critical role of variants involved in gene regulation to promote short-term response to adaptive constraints during invasion. Also, two genes *RhoGEF64C* and *cpo* contained SNPs that were found to be highly significantly associated with the invasive status in both the European and American invasion routes. While the function of the *RhoGEF64C* gene has so far not been extensively studied, several functional and phenotypic studies in other Drosophila species identified genetic variation in the *cpo* gene associated with traits possibly important for invasion success. For instance, *cpo* genetic variation was found to contribute to natural variation in diapause in *D. melanogaster* populations of a North American cline and in populations from the more distantly related species *Drosophila montana* (Kankare *et al.*, 2010; Schmidt *et al.*, 2008). Moreover, indirect action of selection on diapause, by means of genetic correlations involving *cpo* genetic variation, was found on numerous other life-history traits in *D. melanogaster* (Schmidt and Paaby, 2008; Schmidt *et al.*, 2005). Specifically, compared to diapausing populations, non-diapausing populations had a shorter development time and higher early fecundity, but also lower rates of larval and adult survival and lower levels of cold resistance.

Both theoretical (Roughgarden, 1971) and experimental (Mueller and Ayala, 1981) evidence show that traits typical for colonization (i.e., the so-called r-traits; Charlesworth, 1994), such as a non-diapausing phenotype, are selected when a population evolves in a new habitat with low densities and low levels of competition. Common garden studies are needed to assess potential differences in key life history traits (including diapause induction and correlated traits) between native and invasive populations of *D. suzukii* and to evaluate to which extent these are related to the identified variants (including those within the *cpo* gene) differentiating the native and invasive populations of this species.

The *C*_2_ statistic we developed in the present study appears particularly well suited to search for association with population-specific binary traits. Apart from the invasive vs. native status we studied in *D. suzukii*, numerous examples can be found where adaptive constraints may be formulated in terms of contrasting binary population features, including individual resistance or sensibility to pathogens or host-defense systems (e.g., Eoche-Bosy *et al.*, 2017), high vs. low altitude adaptation (e.g., Foll *et al.*, 2014), ecotypes of origin (e.g., Roesti *et al.*, 2015; Westram *et al.*, 2014), or domesticated vs. wild status (e.g., Alberto *et al.*, 2018). In our simulation study, the performance of the *C*_2_ statistic was similar to that of a standard BF obtained after assuming a linear relationship between the (standardized) population allele frequencies and their corresponding binary status. It is worth stressing, however, that *C*_2_ has several critical advantages over BF, as well as over any other decision criterion that may be derived from a parametric modeling.

From a practical point of view, the *C*_2_ estimation does not require inclusion of any other model parameters making it more robust when dealing with data sets including a small number of populations (e.g., <8 populations), the later type of data sets often leading to unstable estimates of BF (unpublished results). In addition, it may easily be derived from only a subset of the populations under study (as we did here when computing the 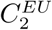 and 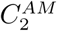 contrasts specific to each of the two invasion routes), while using the complete design to capture more accurate information about the shared population history. Last, the *χ*^2^ calibration of the *C*_2_ under the null hypothesis represents an attractive property in the context of large data sets since it allows to deal with multiple testing issues by controlling for FDR (Francois *et al.*, 2016), via, e.g., the estimation of q–values (Storey and Tibshirani, 2003).

To estimate the *C*_2_ statistic, we needed to correct allele frequencies for population structure. To that end, we relied on the Bayesian hierarchical model implemented in the software BayPass that has several valuable properties including (i) the accurate estimation of the scaled covariance matrix of population allele frequencies (**Ω**), (ii) the integration over the uncertainty of the across population allele frequencies (*π* parameter), and (iii) the inclusion of additional layers of complexities such as the sampling of reads from (unobserved) allele counts in Pool-Seq data (Gautier, 2015). A cost of Bayesian hierarchical modeling is however to shrink the posterior means of the model parameters and related statistics such as the *C*_2_ and XtX differentiation statistics (Gelman *et al.*, 2003). To ensure proper calibration of the two corresponding estimates, we then needed to rely on the rescaled posterior means of the standardized allele frequencies. This empirical procedure proved efficient in providing well behaved p–values while avoiding computationally intensive calibration procedure based on the analysis of pseudo-observed data sets simulated under the generative model (Gautier, 2015). Still, this did not allow accounting for the uncertainty of the allele frequencies estimation (i.e., their full marginal distribution) and more importantly, it implicitly assumes exchangeability of SNPs both across the populations and along the genome. Such an assumption, which pertains to the null hypothesis of neutral differentiation only (and consequently of no association with binary population–specific covariable), might actually be viewed as conservative even in the presence of background LD across the populations, providing that a reasonably large number of SNPs is analyzed. Interestingly, the almost absence of clustering of associated SNPs we observed in the *D. suzukii* genome suggested a very limited extent of across-population LD, presumably resulting from large effective population sizes. This conversely led to a high mapping resolution. In practice, when dealing with large data sets, a sub-sampling strategy consisting in analyzing data sets thinned by marker position also allows further reduction of across-population LD (Gautier *et al.*, 2018). Finally, it should be noticed that information from LD might be at least partially recovered by combining *C*_2_ or XtX derived p–values into local scores (Fariello *et al.*, 2017).

Other less computationally intensive (but less flexible and versatile) approaches may be considered to estimate the *C*_2_ statistic. For instance, the *C*_2_ statistic is closely related to the *S*_*B*_ statistic recently proposed by Refoyo-Martinez *et al.* (2019) to identify footprints of selection in admixture graphs. However, while the *C*_2_ statistic relies on the full scaled covariance matrix of population allele frequencies (**Ω**), the *S*_*B*_ statistic relies on a covariance matrix called ***F*** (Refoyo-Martinez *et al.*, 2019) that specifies an a priori inferred admixture graph summarizing the history of the sampled populations. The covariance matrix ***F*** thus represents a simplified version of **Ω** that may only partially capture the covariance structure of the population allele frequencies. In addition, to compute *S*_*B*_, the graph root allele frequencies are estimated as the average of allele frequencies across the sampled population, which might result in biased estimates, particularly when the graph is unbalanced. Deriving the matrix ***F*** from **Ω** (e.g., Pickrell and Pritchard, 2012) might actually allow interpreting *C*_2_ as a Bayesian counterpart of the *S*_*B*_ statistic, thereby benefiting from the aforementioned advantages regarding the estimation of the parameters Ω and *π* and allowing proper analysis of Pool-Seq data.

### Conclusion and perspectives

Our genome-wide association approach allowed identifying genomic regions and genes most likely involved in adaptive processes underlying the invasion success of *D. suzukii*. The approach can be transposed to any other invasive species, and more generally to any species models for which binary traits of interest can be defined at the population level. The major advantage of our approach is that it does not require a preliminary, often extremely laborious, phenotypic characterization of the populations considered (for example using common garden experiments) in order to inform candidate traits for which genomic associations are sought. As a matter of fact, in our association study the populations analyzed are simply classified into two categories: invasive or native.

The functional and phenotypic interpretation of the signals obtained by our genome scan methods remains challenging. Such interpretation requires a good functional characterization of the genome of the studied species or, failing that, of a closely related species (i.e. *D. melanogaster* in our study). Following a strategy sometimes referred to as “reverse ecology” since it goes from gene(s) to phenotype(s) (Li *et al.*, 2008), it is then necessary to test and validate via quantitative genetic experiments whether the inferred candidate traits show significant differences between native and invasive populations. The functional interpretation of the statistical association results can also benefit from experimental validation approaches based on techniques using RNA interference (RNA-silencing, e.g. Janitz *et al.*, 2006) and/or more genome editing approaches (e.g., Karageorgi *et al.*, 2017) targeting the identified candidate variants. Hopefully, such a combination of statistical, molecular and quantitative approaches will provide useful insights into the genomic and phenotypic responses to invasion, and by the same, will help better predict the conditions under which invasiveness can be enhanced or suppressed

## Materials and Methods

### Simulation study

We used computer simulations to evaluate the performance of the novel statistical framework described in the section *New Approach*. Simulated data sets were generated under the SIMUPOP environment (Peng and Kimmel, 2005) using individual–based forward–in–time simulations implemented on a modified version of the code developed by de Villemereuil *et al.* (2014) for the so-called *HsIMM-C* demographic scenario. This corresponded to an highly structured isolation with migration demographic model (Figure 1A) that was divided in two successive periods: (i) a neutral divergence phase leading to the differentiation of an ancestral population into 16 populations after four successive fission events (at generations *t* = 50, *t* = 150, *t* = 200 and *t* = 300); and (ii) an adaptive phase (lasting 200 generations) during which individuals of the 16 populations were subjected to selective pressures exerted by two environmental constraints (*ec1* and *ec2*), each constraint having two possible modalities (*a* or *b*). We thus had a total of four possible environments in our simulation setting (Figure 1A).

All the simulated populations consisted of 500 diploid individuals reproducing under random-mating with non-overlapping generations. From generation *t* = 150 (with four populations), the migration rate *m*_*jj′*_ between two populations *j* and *j*′ was set to 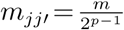 where *p* is the number of populations in the path connecting *k* to *k*′ in the population tree. The migration rate between the two ancestral populations from generation *t* = 50 to *t* = 150 was set to *m* = 0.005. For illustration purposes, some of the migration edges were displayed in Figure 1A.

Following de Villemereuil *et al.* (2014), a simulated genotyping data set consisted of 320 individuals (20 per populations) that were genotyped for 5,000 bi-allelic SNPs regularly spread along ten chromosomes of one Morgan length and with a frequency of 0.5 for the reference allele (randomly chosen) in the root population. Polygenic selection acting during the adaptive phase was simulated by choosing 50 randomly distributed SNPs (among the previous 5,000 ones) that influenced individual fitness according to either the *ec1* or *ec2* environmental constraints (with 25 SNPs for *ec1* and 25 SNPs for *ec2*).

The fitness of each individual, given its genotype, can be defined at each generation. let *p*(*o*)= *j* (*j* = 1,…,16) denote the population of origin of individual *o* (*o* = 1,…,16×500), and *e*_*k*_(*j*)= 1 (respectively *e*_*k*_(*j*)= −1) if the environmental constraint *eck* (*k* = 1,2) of population *j* is of type *a* (respectively *b*). Let further denote *s*_*i*_(*k*) the local selective coefficient of SNP *i* such that *s*_*i*_(*k*)= 0 if the SNP is neutral with respect to *eck* and *s*_*i*_(*k*)= 0.01 otherwise. The fitness of each individual *o* (at each generation) given its genotypes at all the SNPs is then defined using a cumulative multiplicative fitness function as:

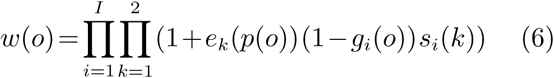

where *g*_*i*_(*o*) is the genotype of individual *o* at marker *i* coded as the number of the reference allele (0, 1 or 2).

### Sampling of *D. suzukii* populations and DNA extraction

Adult *D. suzukii* flies were sampled in the field at a total of 22 localities (hereafter termed sample sites) distributed throughout most of the native and invasive range of the species (Fig 1 and Table S1). Samples were collected between 2013 and 2016 using baited traps (with a vinegar-alcohol-sugar mixture) and sweep nets, and stored in ethanol. Only four of the 22 samples were composed of flies which directly emerged in the lab from fruits collected in the field (Table S1). Native Asian samples consisted of a total of six sample sites including four Chinese and two Japanese localities. Samples from the invasive range were collected in Hawaii (1 sample site), Continental US (6 sites), Brazil (1 site), Europe (7 sites) and the French island of La Réunion (1 site). The continental US (plus Brazil) and European (plus La Réunion Island) populations are representative of two separate invasion routes (the American and European routes, respectively), with different native source populations and multiple introduction events in both invaded areas (Fraimout et al. 2017; see Table S1).

### Pool sequencing

For each of the 22 sampling sites, the thoraxes of 50 to 100 representative adult flies (Table S1) were pooled for DNA extraction using the EZ-10 spin column genomic DNA minipreps kit (Bio basic inc.). Barcoded DNA PE libraries with insert size of ca. 550 bp were further prepared using the Illumina Truseq DNA Library Preparation Kit following manufacturer protocols using the 22 DNA pools samples. The DNA libraries were then validated on a DNA1000 chip on a Fragment Analyzer (Agilent) to determine size and quantified by qPCR using the Kapa library quantification kit to determine concentration. The cluster generation process was performed on cBot (Illumina) using the Paired-End Clustering kit (Illumina). Each pool DNA library was further paired-end sequenced on a HiSeq 2500 (Illumina) using the Sequence by Synthesis technique (providing 2×125 bp reads, respectively) with base calling achieved by the RTA software (Illumina). The Pool-Seq data were deposited in the Sequence Read Archive (SRA) repository under the BioProject accession number PRJNA576997.

Raw paired-end reads were filtered using *fastp* 0.19.4 (Chen *et al.*, 2018) run with default options to remove contaminant adapter sequences and trim for poor quality bases (i.e., with a phred-quality score < 15). Read pairs with either one read with a proportion of low quality bases over 40% or containing more than 5 N bases were removed. Filtered reads were then mapped onto the newly released WT3-2.0 *D. suzukii* genome assembly (Paris *et al.*, 2019), using default options of the *mem* program from the *bwa* 0.7.17 software (Li, 2013; Li and Durbin, 2009). Read alignments with a mapping quality Phred-score < 20 or PCR duplicates were further removed using the *view* (option −q 20) and *markdup* programs from the *SAMtools* 1.9 software (Li *et al.*, 2009), respectively.

Variant calling was then performed on the resulting *mpileup* file using *VarScan mpileup2cns* v2.3.4 (Koboldt *et al.*, 2012) (options *–min– coverage* 50 *–min–avg–qual* 20 *–min-var-freq* 0.001 *–variants-output-vcf* 1). The resulting *vcf* file was processed with the *vcf2pooldata* function from the R package *poolfstats* v1.1 (Hivert *et al.*, 2018) retaining only bi-allelic SNPs covered by > 4 reads, < 99.9th overall coverage percentile in each pool and with an overall MAF> 0.01 (computed from read counts). In total, n=11,564,472 SNPs (respectively n=1,966,184 SNPs) SNPs mapping to the autosomal contigs (respectively X– chromosome contigs) were used for genome-wide association analysis. The median coverage per pool ranged from 58X to 88X and from 34X to 84X for autosomal and X chromosomes, respectively (Table S2). As previously described (Gautier *et al.*, 2018), the autosomal and X-chromosome data sets were divided into sub-data sets of ca. 75,000 SNPs each (by taking one SNP every 154 SNPs and one SNPs every 26 SNPs along the underlying autosomal and X–chromosome contigs, respectively).

### Genome scan analyses

All genome-wide scans were performed using an upgraded version (2.2) of BayPass (Gautier, 2015) (available from http://www1.montpellier.inra.fr/CBGP/software/baypass/), that includes the new *C*_2_ and *XtX* statistics estimated as described in the above section *New Approach*. We always used the BayPass core model with default options for the MCMC algorithm to obtain estimates of four statistical items: (i) the scaled covariance matrix (**Ω**); (ii) the SNP-specific XtX overall differentiation statistic in the form of both 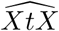, the posterior mean of *XtX* (Gautier, 2015) and 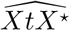, our newly described calibrated estimator; (iii) our novel *C*_2_ statistic in the form of the calibrated estimator described above; and (iv) Bayes Factor reported in deciban units (db) as a measure of support for association with contrasts of each SNP based on a linear regression model (Coop *et al.*, 2010; Gautier, 2015). For BF, a value > 15 db (respectively > 20 db) provides very strong (respectively decisive) evidence in favor of association according to the Jeffreys’ rule (Jeffreys, 1961).

For the *D. suzukii* data sets, we specified the pool haploid sample sizes, for either autosomes or the X–chromosome (Table S1), to activate the Pool-Seq mode of BayPass. The 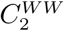 statistic for the contrast of the six native and 16 worldwide invasive populations was estimated jointly with the 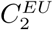 and 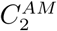 statistics for the contrast of the six native and eight invasive populations of the European and American invasion routes, respectively. For these two latter estimates, this simply amounted to setting *c*_*j*_ = 0 (see eq. 3) for all population *j* not considered in the corresponding contrast analysis. Finally, two additional independent runs (using the option - seed) were performed to assess reproducibility of the MCMC estimates. We found a fairly high correlation across the different independent runs (Pearson’s *r*> 0.92 for autosomal and *r*> 0.87 and X–chromosome data) for the different estimators and thus only presented results from the first run. Similarly and for each chromosome type (i.e., autosomes or the X chromosome), a near perfect correlation of the posterior means of the estimated **Ω** matrix elements was observed across independent runs as well as within each run across SNP sub-samples, with the corresponding FMD distances (Gautier, 2015) being always smaller than 0.4. We thus only reported results regarding the **Ω** matrix that were obtained from a single randomly chosen sub-data set analysed in the first run.

## Acknowledgments

AE, MG and LO acknowledge financial support from the National Research Fund ANR (France) through the project ANR-16-CE02-0015-01 (SWING), the Languedoc-Roussillon region (France) through the European Union program FEFER FSE IEJ 2014-2020 (project CPADROL) and the INRA scientific department SPE (AAP-SPE 2016 and 2018). MGX acknowledges financial support from France Génomique National infrastructure, funded as part of “Investissement davenir” program managed by Agence Nationale pour la Recherche (contract ANR-10-INBS-09). We are grateful to the genotoul bioinformatics platform Toulouse Midi-Pyrenees for providing computing resources, Nicolas Rode for useful discussions and comments on a previous version of the manuscript and Nicolas Ris, Jon Koch, Masahito Kimura, Simon Fellous, Vincent Debat, Marta Pascual, Ruth Hufbauer, Marindia Depra, Isabel Martinez, Pierre Girod and Maxi Richmond for help in collecting some of the *D. suzukii* samples.

---

**TAB. S1:** Description of the 22 *D. suzukii* population samples. The populations representative of the European and American invasion routes are denoted Invasive (EU) and Invasive (AM), respectively (column 3). For each population sample, the thoraxes of 50 to 100 adult flies were pooled; hence the haploid sample sizes of autosomal loci ranging from 100 to 200 (column 7). Pool-samples included both females and male adults, with different proportions of the two sexes depending on the sample; hence the variable number of haploid sample sizes for the X chromosome (column 7).

| Sample name | Sampling site (Lat.; Long.) | Status | Sampling date | Sampling method | Auto. (X) haploid sample size | Auto. (X) median coverage |
| --- | --- | --- | --- | --- | --- | --- |
| CN-Bei | Beijing, China (40.00;116.4) | Native | June 2014 | Baited trap | 100 (89) | 88 (84) |
| CN-Lia | Liaoyuan, China (42.96;125.1) | Native | Aug. 2014 | Baited trap | 100 (42) | 63 (41) |
| CN-Nin | Ningbo, China (30.02;121.5) | Native | July 2014 & May 2016 | Baited trap | 100 (86) | 59 (56) |
| CN-Shi | Shiping county, China (23.7;102.5) | Native | June 2014 & May 2016 | Baited trap | 100 (53) | 61 (34) |
| JP-Sap | Sapporo, Japan (43.05;141.4) | Native | July 2014 | Mullberry | 100 (54) | 77 (40) |
| JP-Tok | Tokyo, Japan (35.64;139.4) | Native | June 2016 | Mullberry, plum | 100 (90) | 62 (56) |
| DE-Dos | Dossenheim, Germany (49.45;8.660) | Invasive (EU) | Aug. 2015 | Baited trap | 100 (58) | 73 (49) |
| DE-Jen | Jena, Germany (50.93;11.56) | Invasive (EU) | Sept. 2016 | Baited trap | 200 (150) | 70 (56) |
| ES-Bar | Barcelona, Spain (41.36;1.964) | Invasive (EU) | July 2014 | Baited trap | 100 (50) | 71 (37) |
| FR-Cor | Corsica, France (42.35;9.003) | Invasive (EU) | Aug. 2016 | Baited trap | 100 (75) | 66 (58) |
| FR-Lez | Montpellier, France (43.70;3.834) | Invasive (EU) | July 2014 | Baited trap | 200 (150) | 82 (65) |
| FR-Par | Paris, France (48.84;2.361) | Invasive (EU) | Nov. 2016 | Baited trap | 200 (150) | 65 (54) |
| FR-Run | La Réunion, France (-21.15;55.64) | Invasive (EU) | Sept. 2016 | Cattley guava | 200 (150) | 86 (68) |
| IT-Tre | Trento, Italy (46.04;11.15) | Invasive (EU) | Sept. 2014 | Baited trap | 200 (140) | 63 (48) |
| BR-Pal | Porto Alegre, Brazil (-27.72;-52.17) | Invasive (AM) | July 2014 | Baited trap | 100 (67) | 68 (53) |
| US-Col | Fort Collins, USA (40.57;-105.1) | Invasive (AM) | Sept. 2015 | Baited trap | 100 (74) | 72 (59) |
| US-Haw | Hawaii (Hilo), USA (19.67;-155.5) | Invasive (AM) | June 2016 | Baited trap | 100 (75) | 87 (71) |
| US-Nca | Raleigh, USA (35.70;-80.62) | Invasive (AM) | Oct. 2016 | Raspberries,Blackberries | 200 (150) | 67 (54) |
| US-Sdi | San-Diego, USA (32.72;-117.2) | Invasive (AM) | May 2014 | Baited trap | 100 (68) | 82 (61) |
| US-Sok | Dayton, USA (45.22;-123.1) | Invasive (AM) | Oct. 2014 | Baited trap, sweep net | 150 (95) | 58 (38) |
| US-Wat | Watsonville, USA (36.90;-121.8) | Invasive (AM) | Oct. 2014 | Sweep net | 100 (54) | 65 (37) |
| US-Wis | Barneveld, USA (42.97;-89.69) | Invasive (AM) | Nov. 2016 | Baited trap | 150 (120) | 70 (58) |

**FIG. S1.**
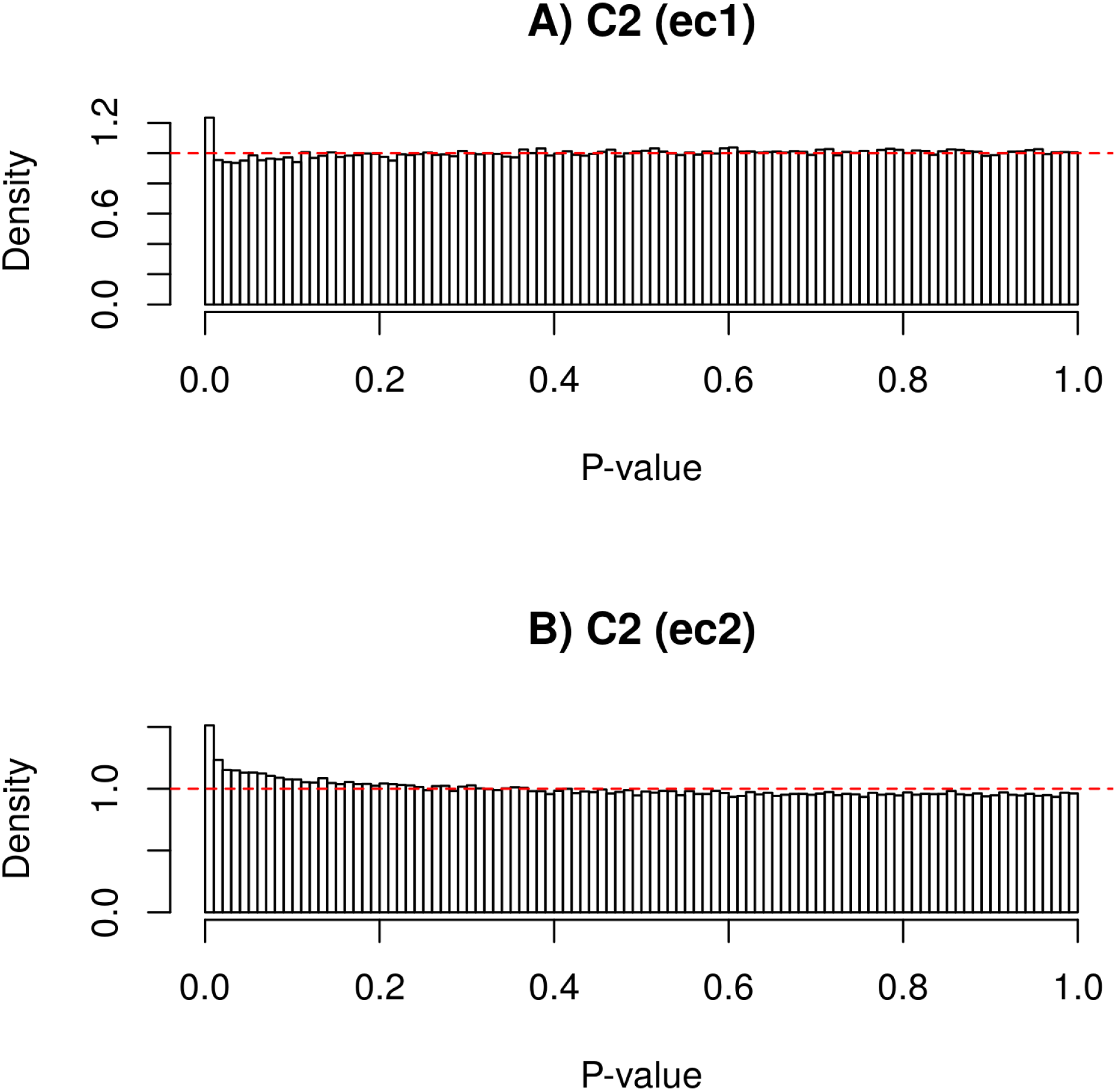
Distribution of the p-values computed on the simulated data (n=500,000 SNPs) and derived from the *C*_2_ statistics for the environmental contrasts *ec1* A) and *ec1* B), assuming a *χ*^2^ null distribution (with one degree of freedom). The red horizontal dashed line represents the uniform distribution.

**FIG. S2.**
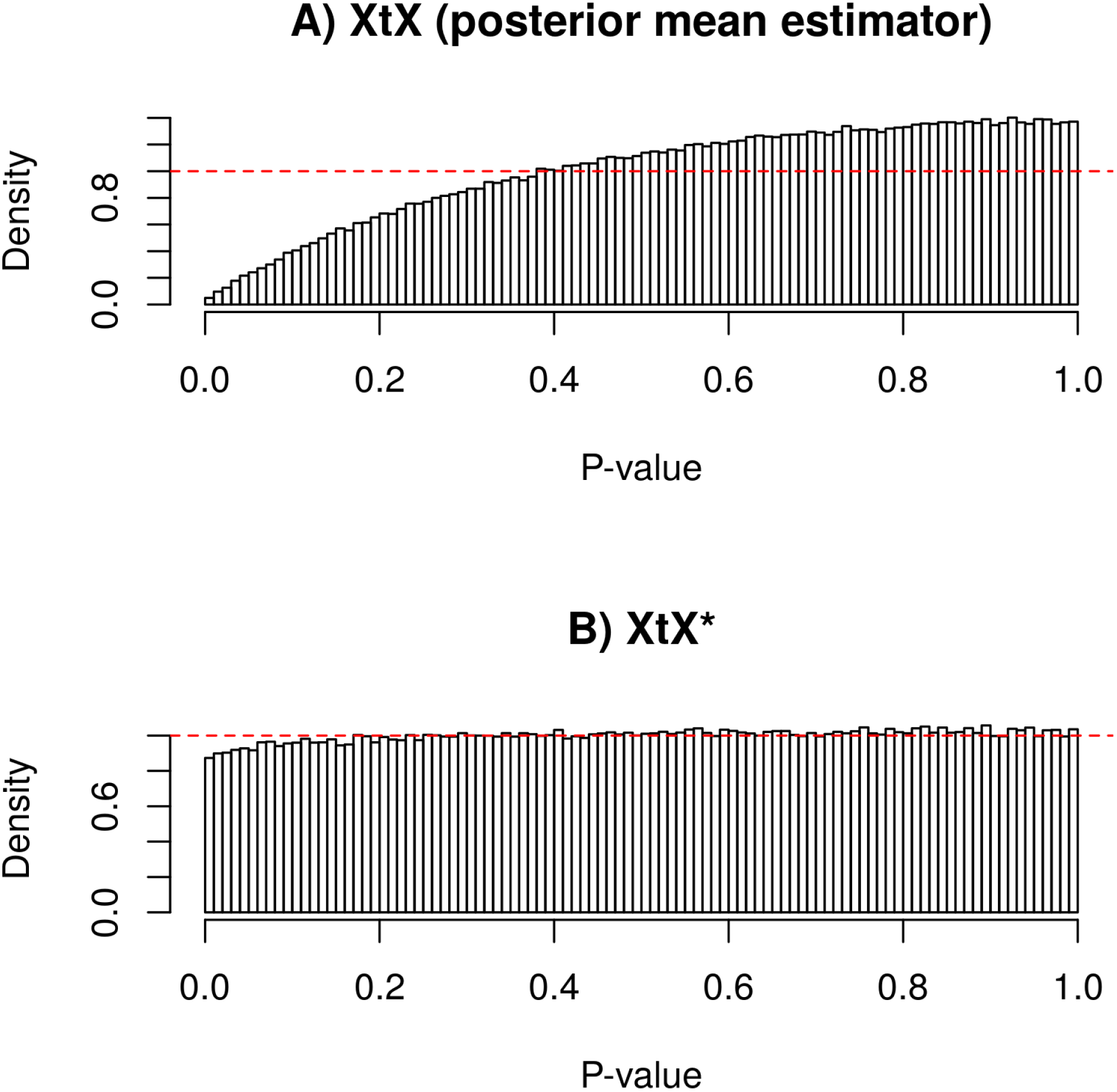
Distribution of the p-values computed on the simulated data sets (n=500,000 SNPs) and derived from the SNP differentiation estimator XtX (posterior mean estimator) A) and the new estimator 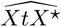 B), assuming a *χ*^2^ null distribution (with *J* = 16, the number of population, degrees of freedom). To account for the bilateral nature of the underlying test (SNPs might be over or under-differentiated if under directional or balancing selection), p-value were computed as 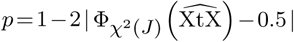, where 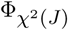 represents the cumulative density function of the *χ*^2^ distribution with *J* degrees of freedom. The red horizontal dashed line represents the uniform distribution.

**FIG. S3.**
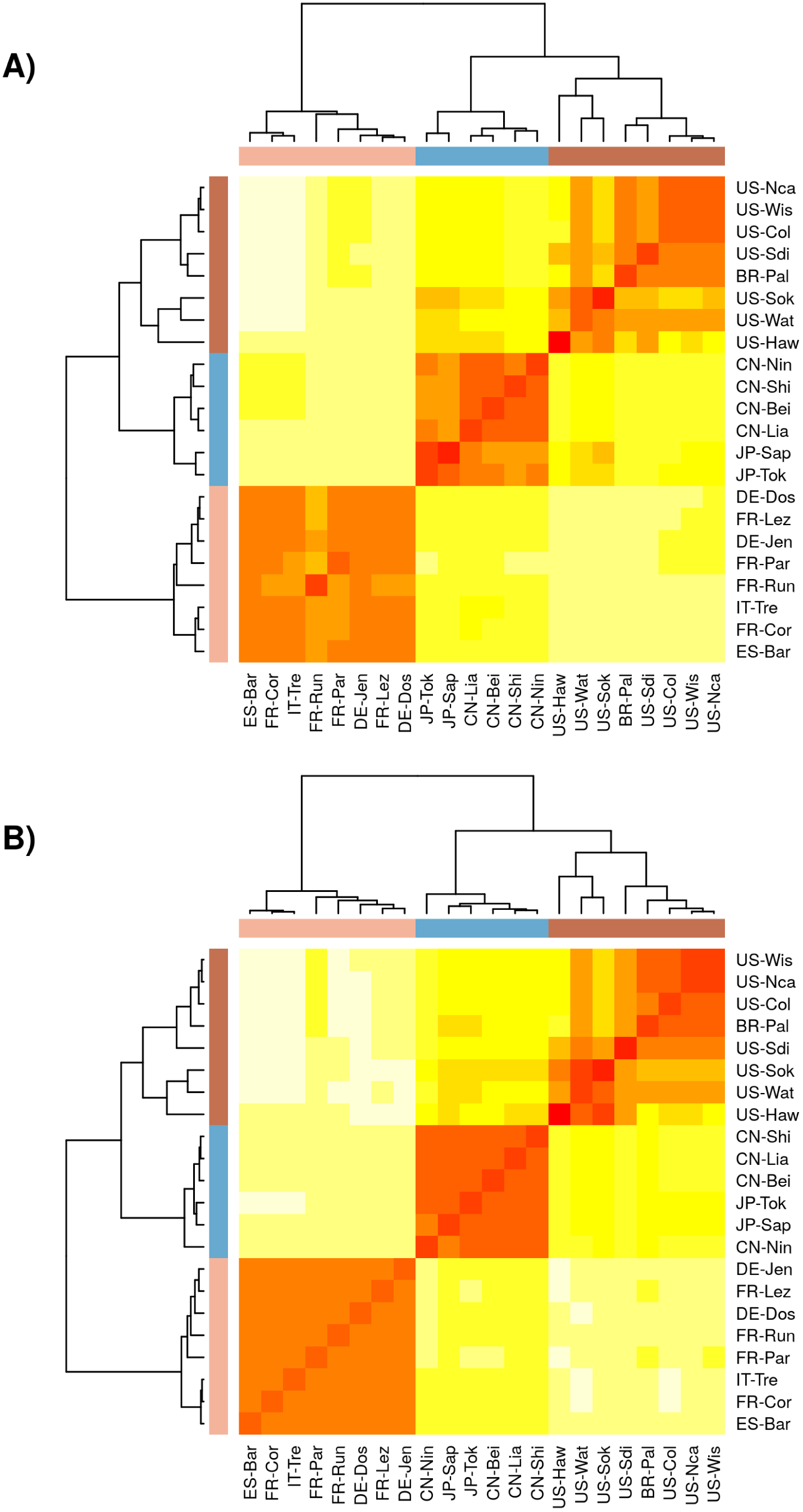
Correlation plot representation of the scaled covariance matrices of population allele frequencies (**Ω**) among all 22 *suzukii* populations based on autosomal (A) and X-linked (B) SNPs.

**FIG. S4.**
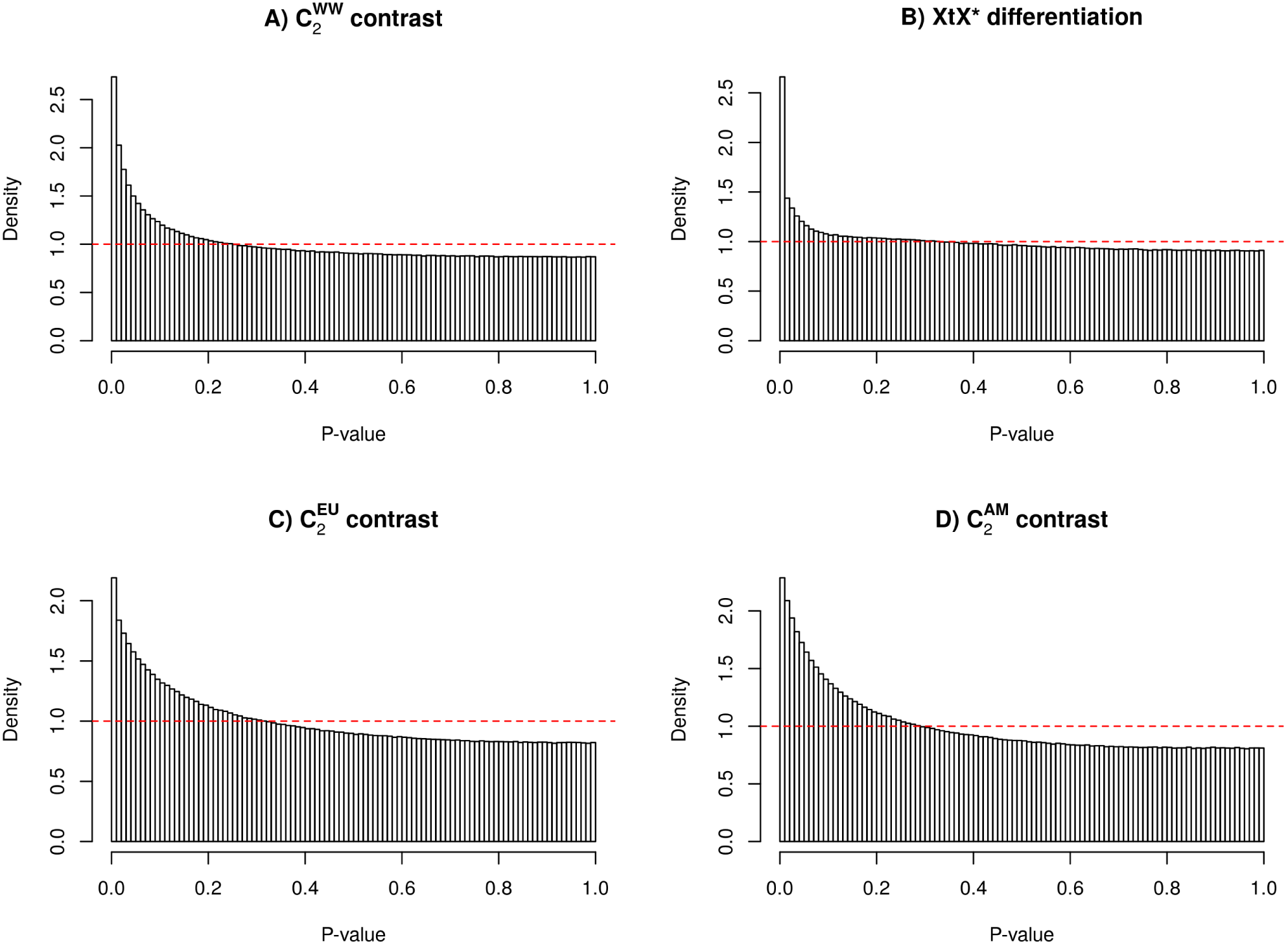
Distribution of the p-values derived from the *C*_2_ contrast statistics estimated from different analyses [A), C) and D)], and from the 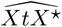 statistic for genetic differentiation among all 22 populations [B)].The *C*_2_ contrast statistics were estimated for 6 native *vs.* 16 worldwide invasive populations 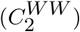 [A)]; 6 native *vs.* 8 invasive populations of the European invasion route 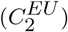 [C)]; and 6 native vs. 8 invasive populations of the American invasion route 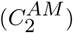 [D)]. The distribution of the (two-sided) p-values derived from the XtX * differentiation statistics (among all 22 *D. suzukii* populations) is given in B). The red dashed line correspond to the uniform distribution.

**FIG. S5.**
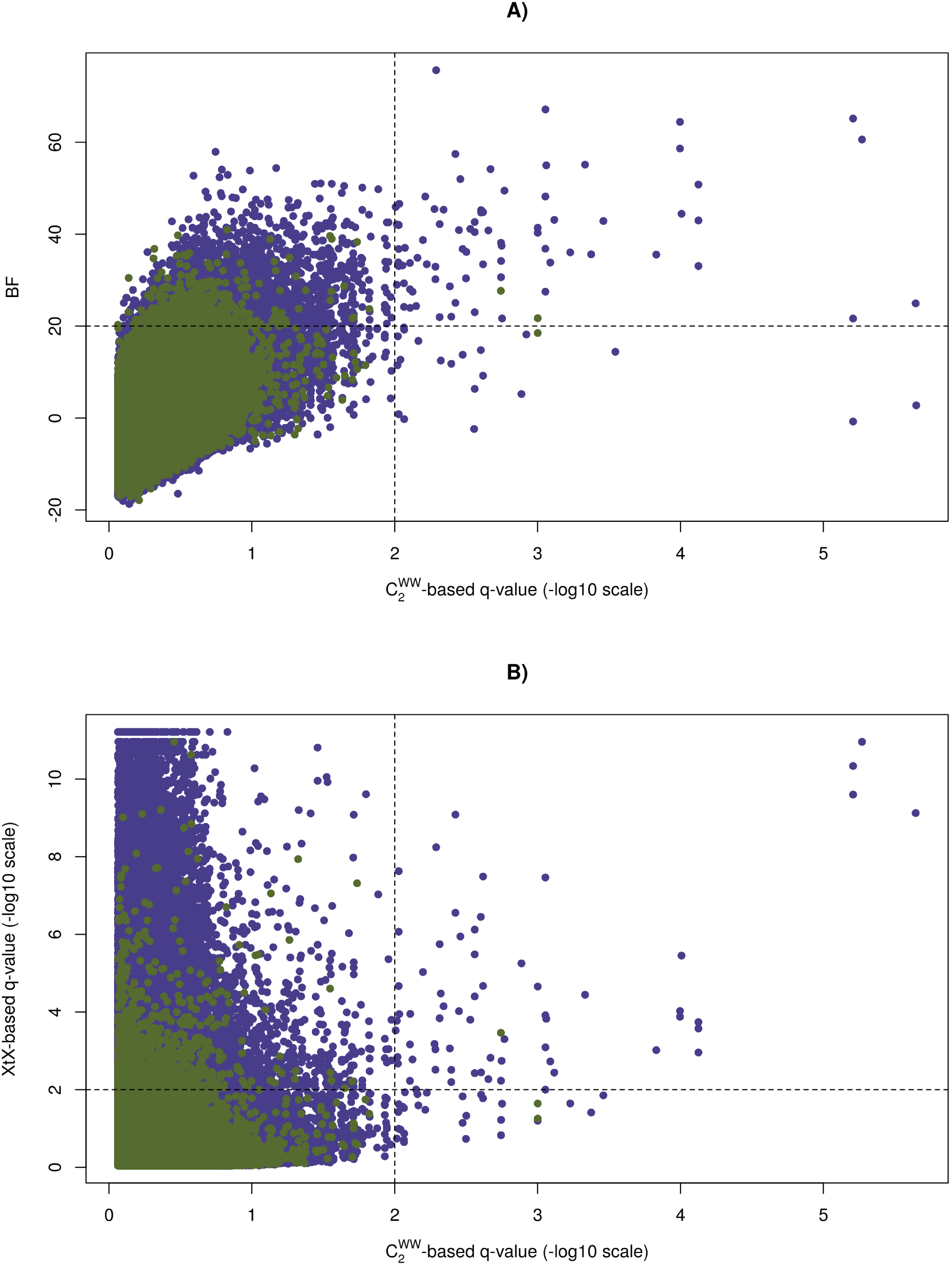
Comparison of the *C*_2_ statistics for the native vs. invasive status of the 22 *D. suzukii* populations 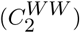 with BF for association A) and with XtX * overall differentiation estimates B). In A) the dashed horizontal line indicates the BF=20 db threshold of decisive evidence according to the Jeffreys’ rule (Jeffreys, 1961) and the dashed vertical line to the 0.1% q–value threshold for the *C*_2_ derived q–values. In B) the horizontal and vertical dashed lines indicate the 0.1% q–value threshold for the XtX * and *C*_2_ derived q–values, respectively. Values for the autosomal (X–linked) SNPs are plotted in purple (green).

**FIG. S6.**
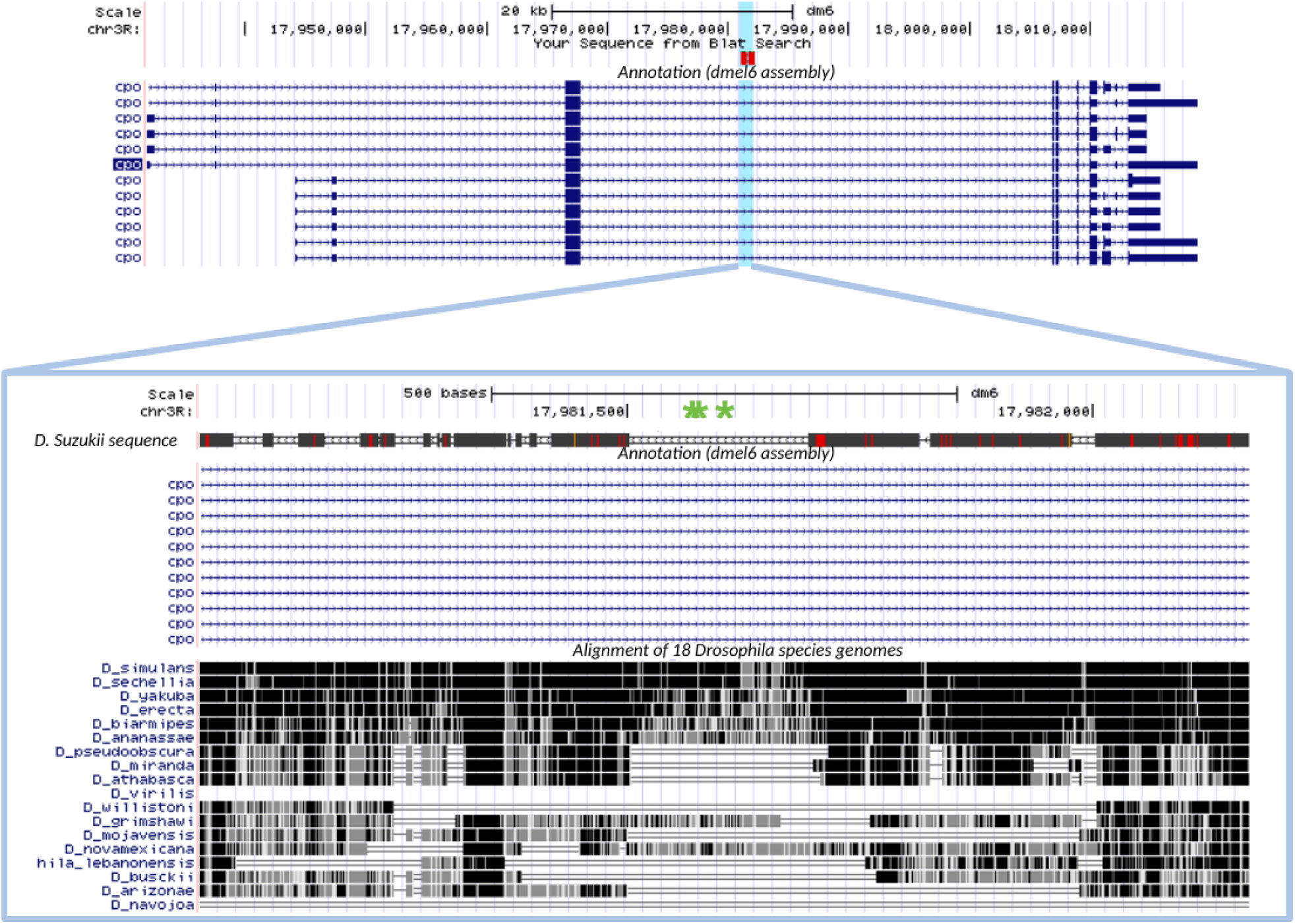
Mapping of the three significant SNPs within the *cpo* gene onto the *dmel6* reference genome of *D. melanogaster* and alignment with genomes from other *drosophila* species. The aligned *D. suzukii* sequence consisted of a 1,193 bp sequence spanning the three significant SNPs (separated by 193 bp) indicated by a green star in the lower panel (the two first SNPs being those significant for the three contrast analyses). The plots were generated using the UCSC genome browser (https://genome.ucsc.edu/).

